## Supplemental File for "An unconventional regulatory circuitry involving Aurora B controls anaphase onset and error-free chromosome segregation in trypanosomes"

//Choose input directory which contains 3D images of metaphase cell.
//An image contains only one metaphase cell. 
//You can have small part of the other cells in the image
//as long as the DAPI (kDNA and Nuc) channel doesn't overlap on your metaphase cell
//This macro analyse all the images in the input directory
input_dir = getDirectory("Input Directory");
list = getFileList(input_dir);

//make index array
row_Index=Array.concat("Image","nuc_vol_um3", "nuc_vol_voxels", "total_protein_int", "KT_int", "KT_voxel");
Array.print(row_Index);

setBatchMode(true);
for (ii=0; ii<list.length; ii++){
	open(input_dir +list[ii]);
	//remove background
	run("Subtract Background...", "rolling=50 stack");
	
	//Clear Results
	run("Clear Results");

	//get original image name
	title=getTitle();
	run("Duplicate...", "duplicate");
	img_copy=getTitle();
	run("Split Channels");

	//set kinetochore and DAPI channels.
	ch_protein=1;
	ch_DNA=2;

	//set shorter names for each images
	img_pro="C"+ch_protein+"-"+img_copy;
	img_DNA="C"+ch_DNA+"-"+img_copy;


	//Select DAPI image
	selectWindow(img_DNA); 

	//find a slice which has the highest intensity pixel, then set it as a focus plane 
	Stack.getStatistics(voxelCount, mean, min, max, stdDev); //get numbers in protein channel
	//go to focus plane, here is the slice which contains highest intensity point (max)
	for (i = 1; i <= nSlices; i++) {
    	setSlice(i);
    	if(max == getValue("Max")){
    		nfocus_DAPI=i;
    	}
	}

	//set threshold to get the default threshold number
	setSlice(nfocus_DAPI);
	setAutoThreshold("Default dark");
	getThreshold(lower, upper);

	//set minimum nuclear size in the image
	getVoxelSize(width, height, depth, unit);
	voxel_size=width * height * depth; //calculate voxel size
	//set min N size to remove kDNA area. 
	//The value came from tne measurement of ~100 cells. DeltaVision images (obj x60)
	minNuc = 2300 * 0.00231125 / voxel_size; 
	
	//set 3D object counter options
	run("3D OC Options", "volume surface nb_of_obj._voxels nb_of_surf._voxels "+
	"integrated_density mean_gray_value std_dev_gray_value median_gray_value "+
	"minimum_gray_value maximum_gray_value centroid dots_size=5 font_size=10 "+
	"white_numbers store_results_within_a_table_named_after_the_image_(macro_friendly) redirect_to=none");
	//get result table and object mask
	run("3D Objects Counter", "threshold="+ lower +" slice=10 min.="+ minNuc +" max.=300560 objects statistics");

	//select measure results
	selectWindow("Statistics for "+img_DNA);

	//empty array to store measurement results
	Measure_results=newArray();
	Measure_results=Array.concat(Measure_results,title);

	//get results from the table and store in Measure_results
	nuc_vol_um3=getResult("Volume (micron^3)", 0);
	Measure_results=Array.concat(Measure_results, nuc_vol_um3);
	nuc_vol_voxels=getResult("Nb of obj. voxels", 0);
	Measure_results=Array.concat(Measure_results, nuc_vol_voxels);
	//close results table
	close("Statistics for "+img_DNA);

	//select 3D mask (nucleus region has intensity 1, other region has intensity 0)
	//apply 3D mask on protein channel
	imageCalculator("Multiply create stack", img_pro, "Objects map of "+img_DNA);

	//select new image
	selectWindow("Result of "+img_pro);
	//find a slice which has the highest intensity pixel, then set it as a focus plane 
	Stack.getStatistics(voxelCount, mean, min, max, stdDev); //get numbers in protein channel
	//go to focus plane, here is the slice which contains highest intensity point (max)
	for (i = 1; i <= nSlices; i++) {
		setSlice(i);
		if(max == getValue("Max")){
    		nfocus_protein=i;
    	}
	}

	//set threshold to get the default threshold number
	setSlice(nfocus_protein);
	setAutoThreshold("Default dark");
	getThreshold(lower, upper);

	//set 3D object counter options
	selectWindow("Result of "+img_pro);
	run("3D OC Options", "volume surface nb_of_obj._voxels nb_of_surf._voxels "+
	"integrated_density mean_gray_value std_dev_gray_value median_gray_value "+
	"minimum_gray_value maximum_gray_value centroid dots_size=5 font_size=10 "+
	"white_numbers store_results_within_a_table_named_after_the_image_(macro_friendly) redirect_to=none");
	run("3D Objects Counter", "threshold="+ lower +" slice=10 min.=1 max.=300560 statistics");

	//get results from the table
	selectWindow("Statistics for Result of "+img_pro);
	No_Obj=Table.size; //number of detected objects
	KT_int=0;
	KT_voxel=0;
	//sum up IntDen for all objects
	for (i = 0; i < No_Obj; i++) {
		KT_int=KT_int + getResult("IntDen", i);
		KT_voxel=KT_voxel + getResult("Nb of obj. voxels", i);
	}
	//print(KT_int);
	close("Statistics for Result of "+img_pro);

	//measure total protein intensity
	selectWindow("Result of "+img_pro);
	//total_int=voxelCount * mean;
	total_int=0;
	run("Set Measurements...", "area mean standard min integrated redirect=None decimal=3");
	for (i = 0; i < nSlices; i++) {
		setSlice(i+1);
		run("Measure");
		total_int=total_int+getResult("RawIntDen", i);
	}
	//close images and results
	close("*");
	close("Results");

	//add results in array
	Measure_results=Array.concat(Measure_results, total_int, KT_int, KT_voxel);
	//print in log window 
	Array.print(Measure_results);
	
}
setBatchMode(false);

selectWindow("Log");
saveAs("text", input_dir+"/Log");
close("Log");
